# Single-chain fluorescent integrators for mapping G-protein-coupled receptor agonists

**DOI:** 10.1101/2023.05.31.543062

**Authors:** Kayla Kroning, Noam Gannot, Xingyu Li, Guanwei Zhou, Jennifer Sescil, Aubrey Putansu, Jiaqi Shen, Avery Wilson, Hailey Fiel, Peng Li, Wenjing Wang

## Abstract

GPCRs transduce the effects of many neuromodulators including dopamine, serotonin, epinephrine, acetylcholine, and opioids. The localization of synthetic or endogenous GPCR agonists impacts their action on specific neuronal pathways. In this paper, we show a series of single-protein chain integrator sensors to determine GPCR agonist localization in the whole brain. We previously engineered integrator sensors for the mu and kappa opioid receptor agonists called M- and K-SPOTIT, respectively. Here, we show a new integrator sensor design platform called SPOTall that we used to engineer sensors for the beta-2-adrenergic receptor (B2AR), the dopamine receptor D1, and the cholinergic receptor muscarinic 2 agonists. For multiplexed imaging of SPOTIT and SPOTall, we engineered a red version of the SPOTIT sensors. Finally, we used M-SPOTIT and B2AR-SPOTall to detect morphine, isoproterenol, and epinephrine in the mouse brain. The SPOTIT and SPOTall sensor design platform can be used to design a variety of GPCR integrator sensors for unbiased agonist detection of many synthetic and endogenous neuromodulators across the whole brain.

## Introduction

G-protein-coupled receptors (GPCRs) are involved in a variety of physiological processes^1^ and are crucial for neuromodulation. GPCRs transduce the effects of many neuromodulators including dopamine, serotonin, epinephrine, acetylcholine, and opioids. Consequently, GPCR mis-regulation is often associated with diseases^2–4^ with over 30% of Federal Drug Administration (FDA)-approved drugs targeting GPCRs^5^. Information on synthetic and endogenous GPCR agonist localization across the whole brain at high spatial resolution is essential for understanding the physiological effects of GPCR synthetic drugs and endogenous GPCR signaling regulation. Brain-wide detection of GPCR agonists is especially important due to the long-range volume transmission of neuromodulators^6^.

To determine the localization of neuromodulators across the whole brain, it is ideal to have a GPCR agonist integrator that can convert the transient agonist signal into a permanent mark on the neurons exposed to the agonists. Permanent labeling of agonist localization has two key advantages over transient labeling: 1. it allows imaging of the agonist localization across the whole brain; 2. the marked neurons can be further analyzed to provide information on the gene expression profile of the neuron. Such genetic information allows the discovery of new druggable targets for treating GPCR mis-regulation.

Various methods have been developed that can provide a real time (or transient) readout in the neurons exposed to GPCR agonists^7–15^. However, fewer tools have been developed that permanently label neurons exposed to GPCR agonists, and the tools that do exist have key limitations. Split fluorescent protein-based^16^ or transcription-based^17–19^ integrators are multi-component systems, where the signal-to-noise ratio (S/N) are highly dependent on the expression level of each protein component. Controlling the relative expression level of the components could be experimentally challenging, requiring extensive optimization^20^. Single protein chain-based integrator systems will address the limitations of multi-component systems.

We previously engineered single protein chain-based integrators called M-SPOTIT (**S**ingle-chain **P**rotein-based **O**pioid **T**ransmission **I**ndicator **T**ool for the **M**u opioid receptor) and its brighter version, M-SPOTIT2 for detecting opioids in cell cultures^21, 22^. In this paper, we show the versatility of the SPOTIT design and describe new single-protein chain fluorescent integrators, including a red-fluorescent SPOTIT, an opioid-activated real-time calcium sensor, and SPOTall, short for **SPOT**IT for **all** GPCRs. We also demonstrate the application of M-SPOTIT for morphine detection and beta-2 adrenergic receptor (B2AR)-SPOTall for isoproterenol and epinephrine detection in animal models. SPOTall provides a new integrator sensor design platform that can be used for the easy engineering of a variety of GPCR agonist integrators. The SPOTIT and SPOTall integrators can be used for whole brain mapping of GPCR agonists.

## Results

### Engineering of SPOTall

The original SPOTIT was designed to specifically detect the agonists for opioid receptors (ORs). To enable the ability to detect the agonists for other GPCRs, here, we established a generalizable sensor design platform that can engineer integrators for many other GPCRs, called SPOTall. We established the SPOTall integrator design platform based on the sensor motif we discovered and utilized in SPOTIT^21^. The sensor motif consists of circularly permuted green fluorescent protein (cpGFP)^23^ and nanobody 39 (Nb39)^24^, where cpGFP fluorophore maturation is inhibited by Nb39 when cpGFP and Nb39 are fused together^21^ and removal of Nb39 results in a >500 times fluorescence increase^21^. This sensor motif was used in the opioid integrator M- SPOTIT, in which cpGFP-Nb39 is attached to the C-terminus of the mu OR (MOR). When agonist binds to M-SPOTIT, Nb39 is recruited to active MOR, removing Nb39 from cpGFP, providing a fluorescent readout (Figure 1a).

**Fig. 1:**
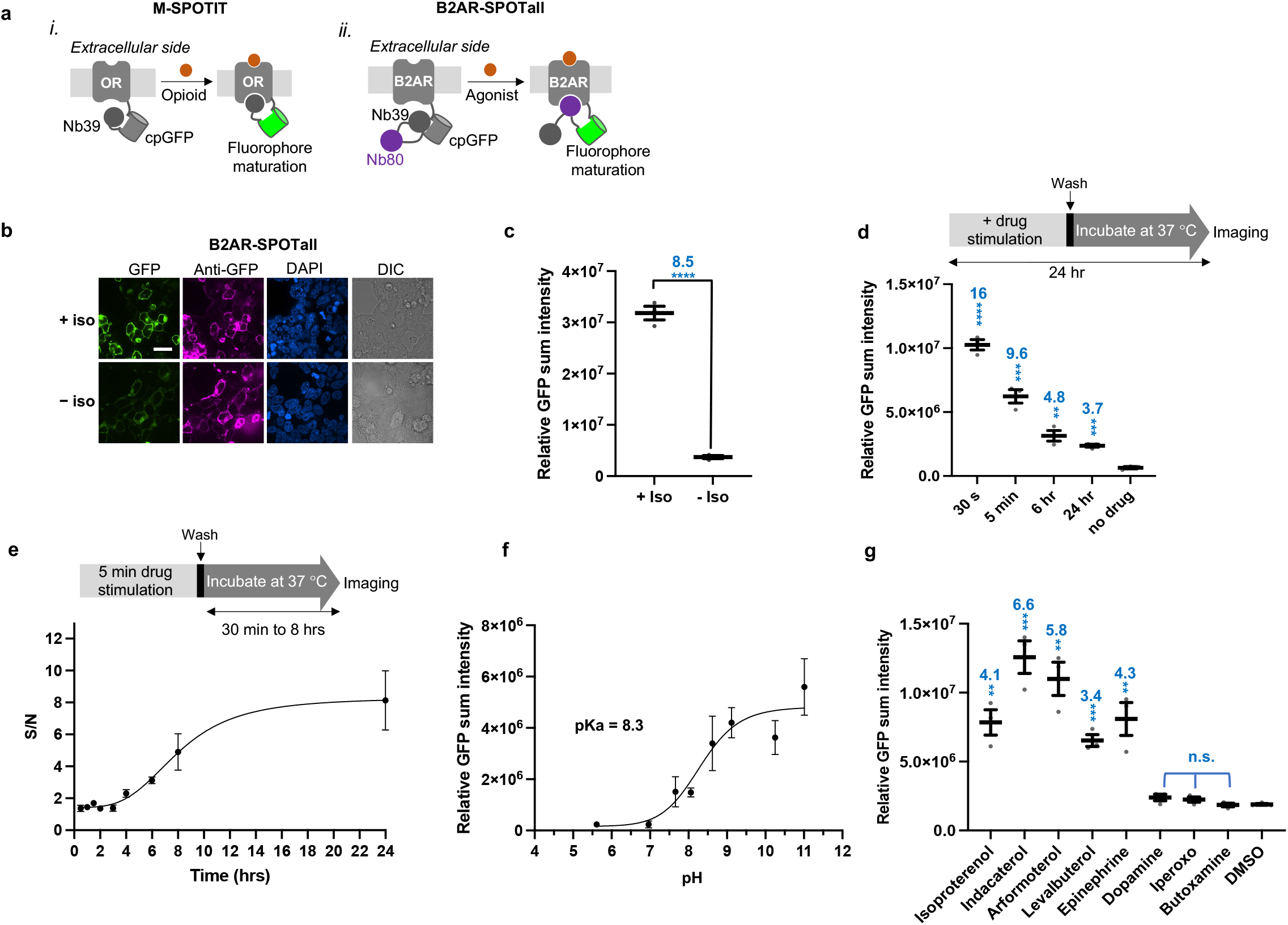
B2AR-SPOTall design and characterizations. **a**, Schematic of M-SPOTIT (*i*) and B2AR-SPOTall (*ii*). For both sensor designs Nb39 inhibits cpGFP fluorophore maturation. For M-SPOTIT, opioid binding recruits Nb39 to the OR, releasing cpGFP and allowing the fluorophore to mature. For B2AR-SPOTall, agonist binding recruits Nb80, sterically blocking Nb39’s interaction with cpGFP and allowing the fluorophore to mature. **b**, Testing B2AR-SPOTall in HEK293T cells. B2AR-SPOTall transfected HEK293T cells were stimulated with 10 µM isoproterenol. 24 hours after stimulation, cells were fixed, immunostained, and imaged with pH 11 buffer. GFP, cpGFP fluorescence. Anti-GFP, protein expression level. DAPI, nuclear staining. DIC, differential interference contrast. Scale bar, 20 µm. Iso, isoproterenol. **c**, Quantification of the experiment described in **b**. The thick horizontal bar is the mean value of three technical replicates. The number above the dots is the S/N; the stars indicate statistical significance. **\*\*\*\***p value < 0.0001. *n*= 3. **d,** Agonist stimulation time testing of B2AR- SPOTall in HEK293T cells. Cells were stimulated with 10 µM isoproterenol for different amounts of time, washed, and then incubated for 24 hours before imaging. Schematic of experiment is shown above the plot. The thick horizontal bar is the mean value of three technical replicates. The number above the dots is the S/N; the stars indicate statistical significance compared to the “no drug” condition. 30 s: ****p< 0.0001, 5 min: ***p= 0.0005, 6 hr: **p= 0.0042, 24 hr: ***p= 0.0003. *n*=3. **e,** Maturation assay of B2AR-SPOTall in HEK293T cells. Cells were imaged 0.5, 1, 1.5, 2, 3, 4, 6, 8, or 24 hours post 5-minute 50 µM isoproterenol stimulation to determine the time it would take the fluorophore to mature. Schematic of experiment is shown above the plot. *n*= 3. Dots on the plot indicate the mean value of three technical replicates. **f**, pH titration of B2AR-SPOTall expressing HEK293T cells. 24 hours post 5-minute 10 µM isoproterenol stimulation, cells were fixed and imaged with different pH buffers: pH 5.6, 7, 7.7, 8.1, 8.6, 9.1, 10.3, and 11. Dots on the plot indicate the mean value of three technical replicates. *n=3.* **g**, Screening B2AR-SPOTall expressing HEK293T cells against different types of ligands. Cells were imaged 24 hours post 5-minute stimulation with 50 µM of drug. n.s., not significant. The thick horizontal bar is the mean value of three technical replicates. The number above the dots is the S/N and the stars indicate significance compared to the “DMSO” condition. Isoproterenol: **p= 0.0030, indacaterol: ***p= 0.0008, arformoterol: **p= 0.0017, levalbuterol: ***p= 0.0004, epinephrine: **p= 0.0067, dopamine: n.s. p= 0.1196, iperoxo: n.s. p= 0.1580, butoxamine: n.s. p= 0.7761. *n= 3.* For c-g, *e*rror bars are the SEM and significance was calculated using an unpaired, two-tailed Student’s *t*-test. A biological replicate has been performed for experiments b-f that yielded similar results.

We harnessed the irreversible fluorophore maturation and large fluorescence dynamic range of the cpGFP-Nb39 sensor motif to engineer SPOTall, an integrator design platform that will be suitable for GPCRs beyond MOR. Since Nb39 is selective for agonist-bound MOR and does not bind to other GPCRs, the SPOTall design requires the incorporation of a conformation-specific binder for the active GPCR of interest. We envisioned a SPOTall design (Figure 1a) in which the conformation-specific binder’s recruitment to the activated GPCR would sterically hinder Nb39’s interaction with cpGFP and allow cpGFP fluorophore maturation.

For the initial testing of the SPOTall design, we chose the B2AR due to its critical function in various physiological processes^25^. Additionally, a conformation-specific nanobody, nanobody 80 (Nb80), already exists with a high binding affinity for activated B2AR^26^. We first tested the control design, in which cpGFP-Nb39 is attached to the C-terminus of B2AR without insertion of Nb80 (Supplementary Figure 1). As expected, the B2AR agonist, isoproterenol did not cause fluorescence increase, because Nb39 is selective towards active ORs. We then tested the B2AR-SPOTall design we envisioned as shown in Figure 1a with Nb80 inserted between cpGFP and Nb39. We also tested another geometry with Nb80 fused after cpGFP-Nb39 (Supplementary Figure 1). B2AR-SPOTall with Nb80 inserted between cpGFP and Nb39 gave significant fluorescence increase upon addition of isoproterenol, while fusing Nb80 after cpGFP-Nb39 did not (Fig.1a-c, Supplementary Figure 1). Therefore, Nb80 inserted in between cpGFP and Nb39 can better disrupt Nb39’s interaction with cpGFP.

With the optimal placement of Nb80, we next probed different linker lengths of the linker connecting B2AR to cpGFP to evaluate its effect on B2AR-SPOTall activation efficiency. We compared the 10 amino acids of the KOR’s C-terminus, as we previously used in M-SPOTIT2^21^, and the truncated linkers with 8 amino acids and 6 amino acids (Supplementary Figure 1). Truncating the linker showed no significant change in the activated signal, so we moved forward with the 10-amino acid linker for B2AR-SPOTall.

### Characterizations of B2AR-SPOTall

We next characterized the various parameters important for B2AR-SPOTall’s performance. The agonist exposure time needed to initiate the sensor activation along with the incubation time needed for fluorophore maturation are both important for SPOTall performance. We previously observed that M-SPOTIT can be activated with a 30-second pulse of fentanyl followed by 12 hours of incubation without fentanyl^21^. Short pulses of agonist create a stable agonist-bound complex that remains after the agonist is washed from the cell media and until the fluorophore has matured.

To characterize the agonist incubation time needed to activate B2AR-SPOTall, we stimulated HEK293T cells expressing B2AR-SPOTall with isoproterenol for 30 seconds, 5 minutes, 6 hours, or 24 hours. After the given stimulation time, the isoproterenol was removed, and the cells were further incubated for 24 hours in a media without agonist to allow the fluorophore to mature. The cells were then fixed and imaged. We found as short as a 30-second stimulation with agonist was sufficient to initiate the B2AR-SPOTall sensor activation (Fig. 1d). However, longer agonist incubation times resulted in decreased fluorescence activation (Fig. 1d), which we hypothesized was from lower protein level due to hyperstimulation of B2AR. For the remaining cellular characterizations of B2AR-SPOTall, we used a 5-minute agonist stimulation protocol, which is easier to perform than the 30-second stimulation and still gives a relatively high fluorescent increase upon agonist stimulation.

To evaluate the time needed for fluorophore maturation in B2AR-SPOTall, we stimulated B2AR-SPOTall expressing cells for 5 minutes followed by further incubation for differing amounts of time before imaging. Clear agonist-dependent activation was observed 8-hours post stimulation, reaching a plateau at around 24 hours (Fig. 1e). Moving forward, we imaged cells 24 hours post stimulation to give the highest sensor activation and S/N. Here, we define the S/N as the ratio of the agonist-stimulated fluorescence to the unstimulated condition. Even though SPOTall requires >8 hours of incubation for fluorophore maturation, the agonist needs to be incubated as short as 30 seconds to form a stable complex to initiate fluorophore maturation.

Another parameter that is important for SPOTall’s performance is the fluorophore’s pKa in a formaldehyde-fixed condition. We observed previously that cpGFP in fixed cells has a high pKa, requiring high pH buffer to deprotonate the fluorophore to observe maximum fluorescence ^21^. We next measured the pKa of the B2AR-SPOTall fluorophore formed in formaldehyde-fixed conditions. We performed a pH titration by imaging the stimulated B2AR-SPOTall with buffers at a range of pH values. We determined the B2AR-SPOTall’s pKa to be 8.3 (Fig. 1f), requiring a pH >10.3 to achieve >99% deprotonation of the fluorophore. Therefore, a buffer at pH 11 is used for B2AR-SPOTall imaging for all experiments.

To evaluate B2AR-SPOTall’s selectivity for B2AR agonists, we characterized B2AR-SPOTall against a variety of ligands, including FDA-approved agonists, the endogenous ligand, an antagonist, and ligands selective for other GPCRs. B2AR-SPOTall was significantly activated by B2AR agonists but not by the antagonist or agonists for other GPCRs (Fig. 1g), validating the selectivity of the B2AR-SPOTall for B2AR agonists.

Lastly, to evaluate B2AR-SPOTall’s sensitivity for B2AR agonists, we stimulated B2AR-SPOTall with ligands of varying concentrations. The FDA-approved drugs arformoterol, indacaterol, and isoproterenol all showed EC50 values of approximately 1-6 μM (Supplementary Fig. 2). B2AR-SPOTall’s EC50 value for epinephrine is 10 times higher (44 μM) compared to the FDA-approved drugs. This data shows that B2AR-SPOTall is more sensitive towards synthetic ligands than the endogenous agonist.

### Easy extension of the SPOTall platform to other GPCRs

To test the generalizability of the SPOTall platform, we next applied the optimal SPOTall design to other GPCRs by simply replacing B2AR and Nb80 with other GPCR-conformation-specific binder pairs. We first engineered a SPOTall sensor with another G^s^-coupled receptor, the dopamine receptor D1 (DRD1). We reasoned DRD1’s structural similarity to B2AR might allow Nb80 to bind to the active DRD1. Therefore, the only change we made to the B2AR-SPOTall design for DRD1-SPOTall is that we replaced B2AR with DRD1 (Fig. 2a). A significant fluorescence increase was observed upon addition of dopamine with a S/N of 43. This shows that Nb80 can bind to activated DRD1 intramolecularly and that the SPOTall sensor design can be extended to another GPCR (Fig. 2b and c).

**Fig. 2:**
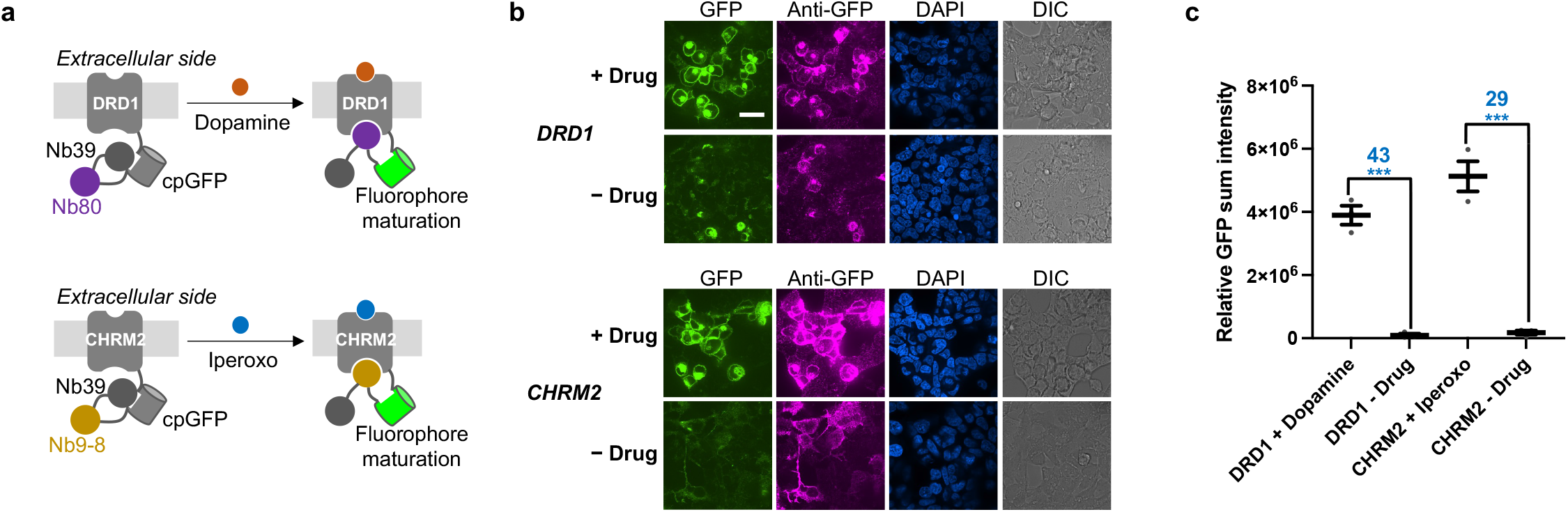
Design and testing of DRD1- and CHRM2-SPOTall. **a**, Schematic of DRD1- and CHRM2-SPOTall. **b**, HEK293T cell testing of DRD1- and CHRM2-SPOTall. HEK293T cells expressing DRD1- and CHRM2-SPOTall were stimulated with 100 µM dopamine and iperoxo, respectively. Cells were fixed, immunostained, and imaged 24 hours post stimulation. GFP, cpGFP fluorescence. Anti-GFP, protein expression level. DAPI, nuclear staining. DIC, differential interference contrast. Scale bar, 20 µm. **c**, Quantification of **b**. Error bars are the SEM. The thick horizontal bar is the mean value of three technical replicates. The number above the dots is S/N and the stars indicate significance compared to the “− drug” condition. Significance was calculated using an unpaired, two-tailed Student’s *t*-test. DRD1: ***p value= 0.0002, CHRM2: ***p value= 0.0005. *n*= 3. A biological replicate has been performed for experiments b and c that yielded similar results.

We next engineered a SPOTall sensor with another GPCR, the cholinergic receptor muscarinic 2 (CHRM2). CHRM2 also has an available conformation-specific nanobody, Nb9-8^27^. We designed CHRM2-SPOTall as shown in Fig. 2a. A significant fluorescence increase was observed upon addition of the CHRM2 agonist iperoxo with a S/N of 29 (Fig. 2b and c). These studies show that the SPOTall platform is generalizable for GPCRs with a conformation-specific binder. However, both DRD1-SPOTall and CHRM2-SPOTall have overall lower fluorescence intensity than B2AR-SPOTall, necessitating further improvement before animal testing.

### Engineering of red-SPOTIT

In addition to extending the single-chain integrator to other GPCRs, we designed a single-chain red fluorescent integrator. A red fluorescent integrator will complement the green fluorescent SPOTIT and SPOTall for multiplexed detection of GPCR agonists. Red fluorescent sensors are also advantageous in that red wavelengths overlap less with the autofluorescence of animal tissue, improving the S/N.

In the red-SPOTIT design, we replaced cpGFP with circularly permuted mApple (cpmApple), a red fluorescent protein used in the calcium sensor, O-Geco1^28^. Surprisingly, this design showed no cpmApple fluorescence with or without opioid stimulation (data not shown). To investigate this, we performed a control study of expressing cpmApple alone in comparison to cpmApple fused to the calcium-dependent protein pair, M13 and CaM (the same construct as O-Geco1). Interestingly, the cpmApple alone from O-Geco1 does not have fluorescent signal at pH 11 while O-Geco1 shows red fluorescence (Supplementary Fig. 3). The lack of fluorescence in cpmApple alone is not due to protein stability since immunofluorescence indicates the protein is expressed for both constructs. This suggests that the cpmApple fluorophore in O-Geco1 requires fusion to M13 and calmodulin to mature. Therefore, to engineer red-SPOTIT, we need to incorporate the whole O-Geco1 construct rather than just cpmApple itself.

To determine if Nb39 can inhibit cpmApple fluorophore maturation with M13 and calmodulin added back, we added Nb39 to the C-terminus of O-Geco1 and saw significant inhibition of fluorophore maturation (Supplementary Fig. 3). Additionally, protease cleavage to remove Nb39 from O-Geco1 can significantly increase the red fluorescence (Supplementary Fig. 3). This suggests that O-Geco1-Nb39 works similarly to cpGFP-Nb39 and can be used for a single-chain fluorescent integrator design.

To design a red opioid integrator, we fused O-Geco1-Nb39 to the C-terminus of the MOR and kappa opioid receptor (KOR) (Fig. 3a). These two red fluorescent integrators are called M- and K-red-SPOTIT. M- and K-red-SPOTIT each yielded a fluorescence increase upon opioid stimulation with a S/N of 4.2 and 4.4, respectively (Fig. 3b and Supplementary Fig. 3).

**Fig. 3:**
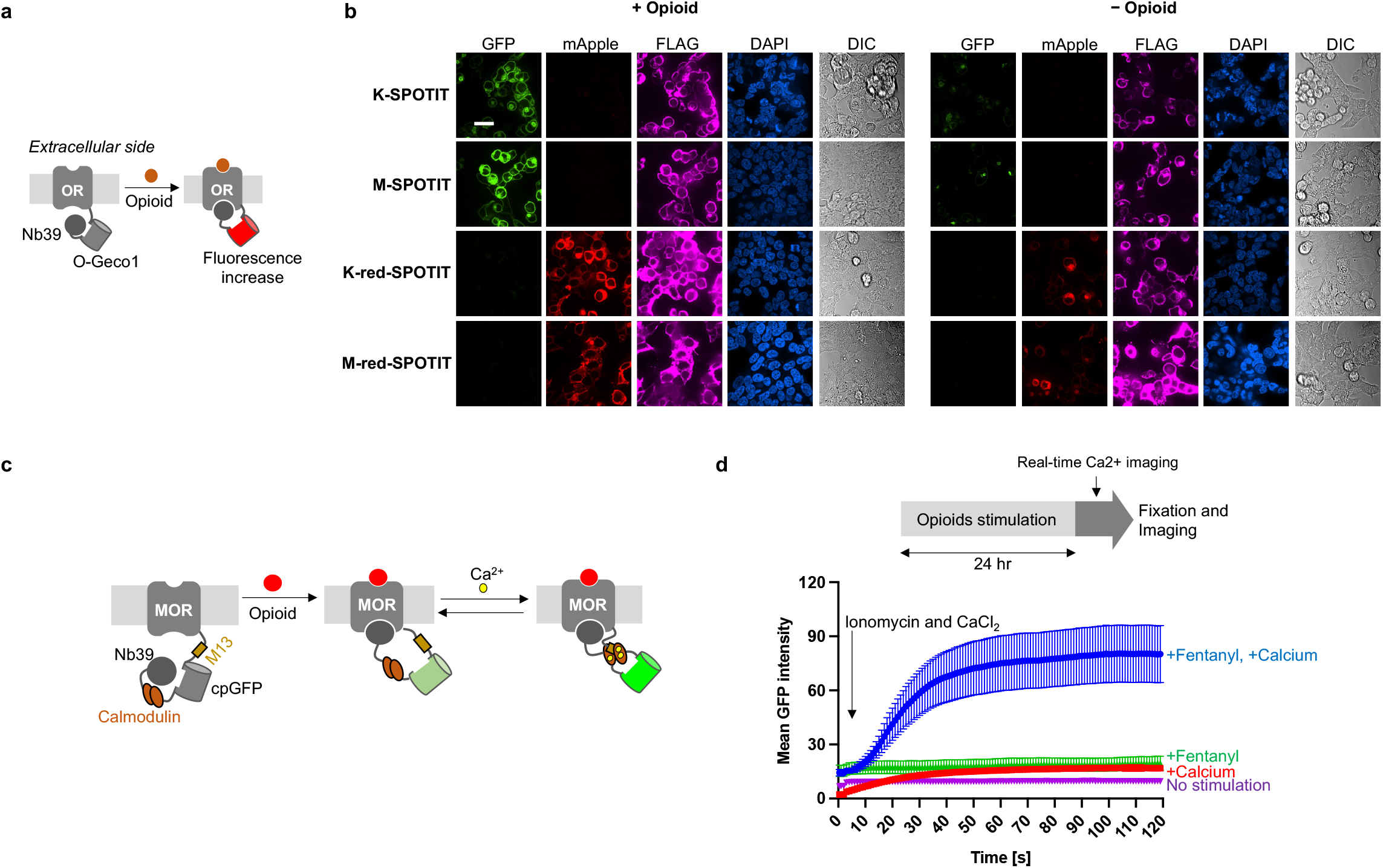
red-SPOTIT and SPOTcal design and testing. **a**, Schematic of red-SPOTIT design. O-Geco1 replaces cpGFP from the original SPOTIT design. Nb39 inhibits O-Geco1 fluorophore maturation. Opioid binding recruits Nb39 to the OR, allowing the O-Geco1 fluorophore to mature. **b**, HEK293T cell testing of green or red SPOTIT. SPOTIT expressing cells were stimulated with 10 µM of fentanyl or salvinorin A for MOR or KOR, respectively. 24 hours post opioid stimulation, cells were fixed, immunostained, and imaged at pH 11. GFP, cpGFP fluorescence. mApple, cpmApple fluorescence. FLAG, protein expression level. DAPI, nuclear staining. DIC, differential interference contrast. Scale bar, 20 µm. **c**, Schematic of the opioid-activated calcium sensor. Opioid binding allows the fluorophore to mature, where calcium binding leads to an increase in fluorescence in live cells. **d,** Schematic of SPOTcal testing and plot of the real-time fluorescence increase from calcium stimulation. HEK293T cells expressing the opioid-activated calcium sensor were stimulated with 10 µM fentanyl 24 hours post transfection. 24 hours after fentanyl stimulation, cells were then imaged in real-time before and after addition of 5 mM calcium chloride and 2 µM ionomycin. Controls were also performed without opioid or without calcium and ionomycin. Mean values of replicates are indicated by the dot and error bars are the SEM. *n=* 30.

### Engineering of an opioid-activated real-time calcium sensor

Calcium influx is a common marker for neural activity. Consequently, many genetically encoded calcium sensors have been engineered to determine neural activity in cell-type specific neural populations^29^. By combining the SPOTIT design and the widely used calcium sensor, GCaMP6^23^, we engineered a real-time calcium sensor that is only functional in opioid-dependent neural circuits. We reasoned such a sensor can be used to detect neuronal activity induced calcium influx specifically in the neuronal circuits involved in opioid signaling, offering more specificity to the GCaMP sensor.

To design an opioid-activated calcium sensor, we replaced cpGFP in green M-SPOTIT with M13-cpGFP-CaM from GCaMP6^22^ (Fig. 3c). In this design, the fluorophore maturation is still dependent on opioid stimulation. Once the integrator’s fluorophore is matured after opioid stimulation, the integrator can then serve as a real-time calcium indicator, because it contains the GCaMP6 elements (Fig. 3c). We call this new integrator SPOTcal.

To test SPOTcal, we stimulated SPOTcal expressing HEK293T cells with fentanyl to induce fluorophore maturation first. We waited 24 hours for significant fluorophore maturation to take place and then tested the matured SPOTcal’s real-time calcium sensing ability. We stimulated the cells with calcium and ionomycin to increase intracellular calcium concentration. Ionomycin and calcium treatment caused a fluorescence increase in both the fentanyl and non-fentanyl treated cells with higher fluorescence in the cells previously stimulated with fentanyl (Fig. 3d and Supplementary Fig. 4), illustrating the opioid-dependent calcium detection of SPOTcal.

### Animal application of M-SPOTIT2 and B2AR-SPOTall

To evaluate the potential of these single-chain fluorescent integrators for detecting GPCR agonists in animal models, we tested M-SPOTIT and B2AR-SPOTall in a mouse brain. We first tested M-SPOTIT2^22^ by injecting home-made adeno associated viruses (AAV) 1/2 mixed serotype encoding M-SPOTIT2 into the preBötzinger complex (preBötC) of a mouse brain. Due to the large injection volume, viral expression occurred in regions beyond the preBötC as well. The preBötC is the inspiratory rhythm generator and is impacted by opioid signaling^30^.

6 days after viral injection of M-SPOTIT2, we administered morphine or saline via IP injection to the mice. Then, 24 hours after morphine or saline administration, we sacrificed the mice for imaging (Fig. 4a). We observed a 4.9 times signal increase in the morphine administered group in comparison to the saline control group (Fig. 4b and c and Supplementary Fig. 5). Lower M-SPOTIT2 protein levels were observed in the saline stimulated condition (Fig. 4b). Lower protein levels in the absence of drug has previously been observed for the SPOTIT sensors^31^ and is due to the instability of the inactive cpGFP-Nb39 complex. The higher signal in the morphine stimulated condition is, therefore, due to both fluorophore maturation and higher sensor stability. This study shows that M-SPOTIT2 can detect synthetic opioids in a mouse brain.

**Fig. 4:**
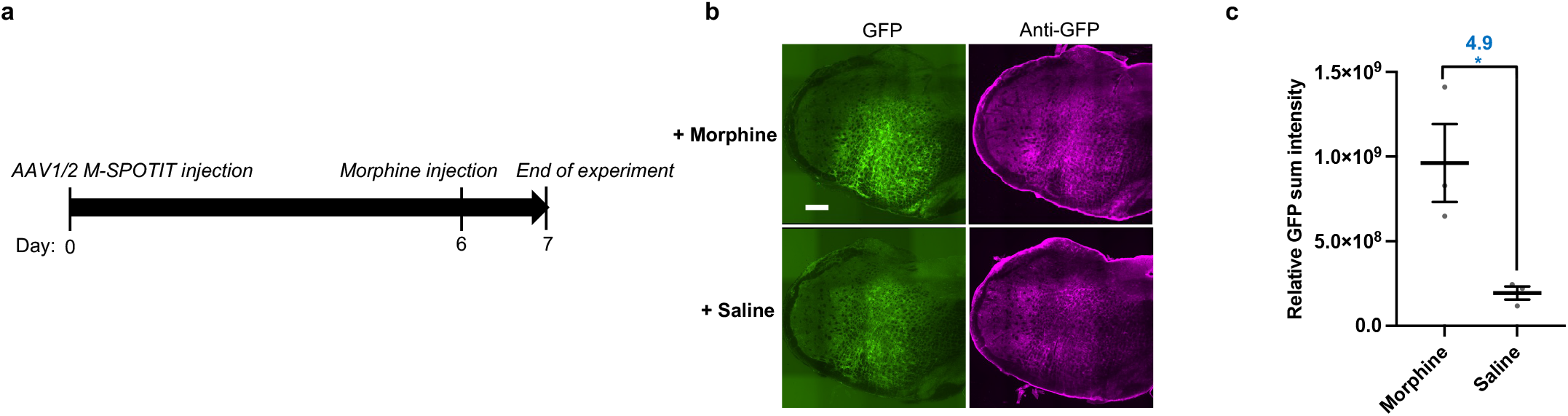
Mouse testing of M-SPOTIT2 with AAV1/2 viral serotype. **a**, Experimental protocol for M-SPOTIT2 mouse testing. **b**, Representative images of M-SPOTIT2 mouse testing. 100 mg/kg of morphine or saline was administered through intraperitoneal injection 6-days after viral delivery to the preBötC. GFP, cpGFP fluorescence. Anti-GFP, protein expression levels. Scale bar, 300 µm. Images cropped to show injection site. Uncropped images are found in Supplementary fig. 5. **c**, Statistics of experiment described in **b**. Mean is represented by the horizontal bar. Each dot is one animal. Error bars, SEM. Stars indicate significance after performing an unpaired, two-tailed Student’s t-test. Morphine: *p= 0.0303. *n*= 3.

It is important to note that the SPOTIT sensors can also engage in G-protein signaling. To characterize M-SPOTIT2’s G-protein signaling abilities, we used the GloSensor^32^ assay to measure cAMP levels in cell cultures. Fentanyl stimulation with M-SPOTIT2 reduces the cAMP levels in HEK293T cells, indicating M-SPOTIT2 can couple to G-proteins (Supplementary Figure 6). To prevent G-protein coupling to M-SPOTIT2, we incorporated a series of mutations in the G-protein binding pocket of MOR^33^. We found one mutant, R279F, that has abolished G-protein coupling while still maintaining sensor activity (Supplementary Fig. 6). These mutants can be used if G-protein binding to M-SPOTIT2 becomes a concern in certain experiments. To illustrate this mutation strategy works for other GPCR sensors, we applied the same strategy to the KOR sensor, K-SPOTIT, where we were also able to reduce G-protein coupling to K-SPOTIT (Supplementary Fig. 7). We reasoned this strategy can be used for all future SPOTall sensors if G-protein coupling is a major concern.

Lastly, we tested the application of B2AR-SPOTall in mouse brain. AAV vector encoding B2AR-SPOTall was injected into the lateral hypothalamus area. Because the activated fluorescence signal and background of B2AR-SPOTall are both lower than M-SPOTIT2 in HEK293T cells (Supplementary Fig. 8), we extended the viral expression time to a total of 7 days before drug or saline administration (Fig. 5a). Both isoproterenol and epinephrine administration induced an increase of green fluorescent signal (6.0- and 3.7-fold, respectively), compared to the saline administered mice (Fig. 5b and c and Supplementary Fig. 9). Therefore, B2AR-SPOTall can be applied in vivo to detect the localization of neuromodulators.

**Fig. 5:**
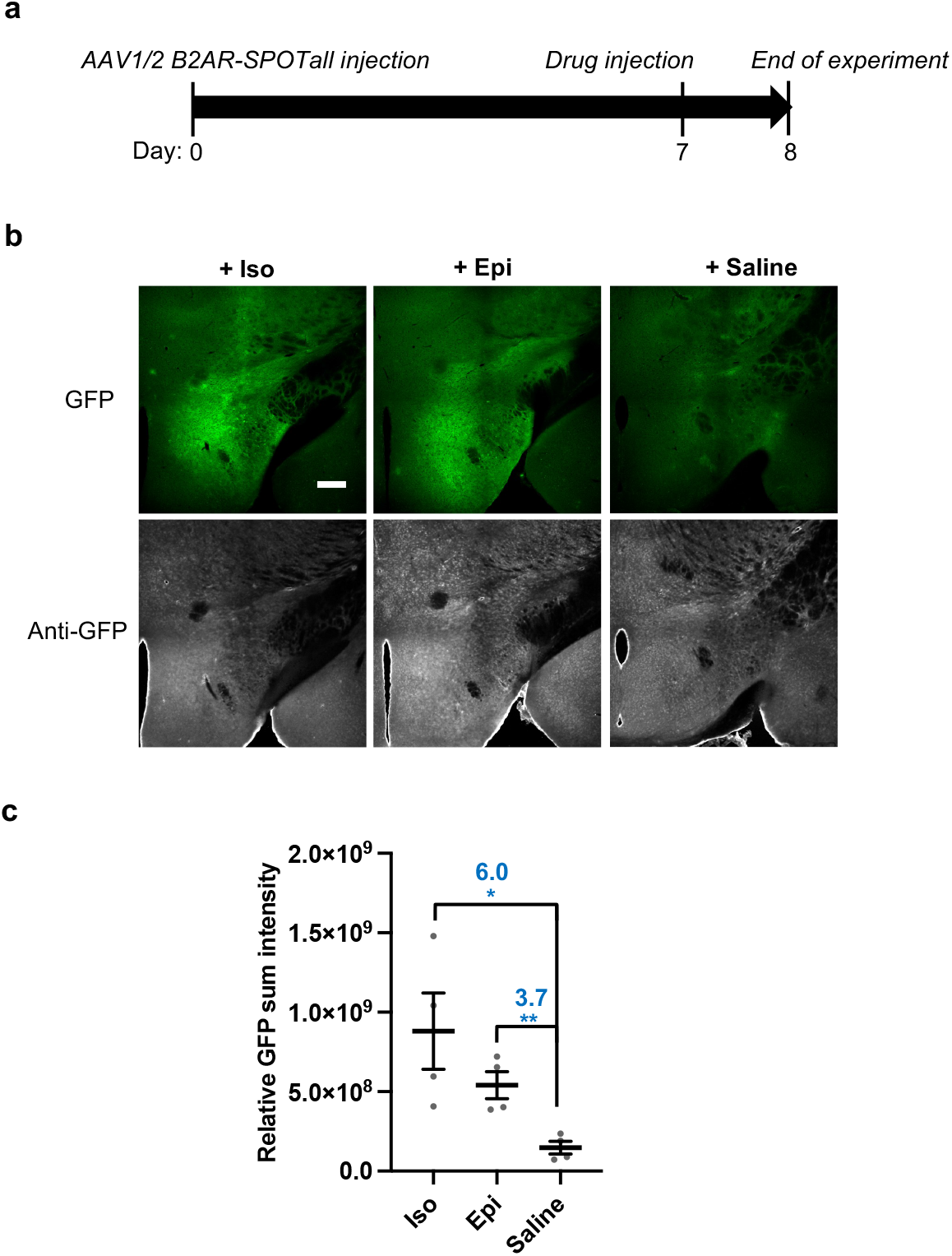
Mouse testing of B2AR-SPOTall with AAV1/2 viral serotype. **a**, Experimental protocol for B2AR-SPOTall mouse testing. **b**, Representative images of mouse testing. 1µL of 10 mM isoproterenol, 10 mM epinephrine, or saline was locally injected to the lateral hypothalamus area 7 days after viral delivery to the same area. GFP, cpGFP fluorescence. Anti-GFP, protein expression levels. Scale bar, 300 µm. **c**, Statistics of experiment described in **a** for M-SPOTIT. Mean is represented by the horizontal bar. Each dot is one animal. Error bars, SEM. Stars indicate significance after performing an unpaired, two-tailed Student’s *t*-test. Iso: *p= 0.0233. Epi: **p= 0.0059. *n*= 3. Iso, isoproterenol. Epi, epinephrine.

## Discussion

Integrators for GPCR agonists are needed for whole-brain mapping of GPCR agonist localization at cellular resolution and allow the analysis of neurons post signaling event, where the genetic profile of the neuron can be determined. SPOTIT and SPOTall do not have time gating; however, time gating can be achieved by adding temporal controls to integrators, such as expressing integrators under a drug inducible promoter^34^.

In this paper, we showed the design of multiple single-chain fluorescent integrators for detecting GPCR agonists. In addition to our previously designed green fluorescent M-SPOTIT for detecting MOR agonists, we designed SPOTall that is more generalizable for a range of GPCRs and a red-SPOTIT for potential multiplexed detection. We demonstrated SPOTall’s easy adaption with three different GPCRs: B2AR, DRD1, and CHRM2. Notably, B2AR-SPOTall can detect as short as 30 seconds of agonist stimulation and has high selectivity towards B2AR agonists. We also designed an opioid-activated calcium sensor, where calcium sensing occurs only in neurons previously exposed to opioids. This sensor offers important specificity to the widely used calcium sensor, GCaMP6^23^. To demonstrate these sensors’ potential applications in mouse models, we also showed the proof-of-principle applications of M-SPOTIT2 and B2AR-SPOTall in mouse brains.

B2AR-SPOTall and M-SPOTIT2 are the first single-chain fluorescent integrators for detecting GPCR agonists. They will be useful for detecting where B2AR- and MOR-targeted drugs are localizing in the brain globally at cellular resolution. After further improving the sensitivity for their endogenous agonists, B2AR-SPOTall and M-SPOTIT2 will also be useful for detecting the release of endogenous B2AR and MOR agonists to better understand the spatial regulation of endogenous signaling.

The SPOTall design is highly modular and can be extended to design integrators for other GPCRs. While nanobodies are used as protein binders for active GPCRs in the current SPOTall design, non-nanobody protein binders can also be tested in the future. Additionally, M-red-SPOTIT can potentially be used at the same time as a SPOTall sensor to observe how opioid signaling impacts other neuromodulator signaling events. For example, evidence has shown that endogenous opioid peptides can modulate epinephrine release^35^ and synthetic opioid agonists can impact dopamine release^36^. Red or green SPOTIT and SPOTall can also be used in combination with real-time sensors to first determine the localization of GPCR agonists throughout the whole brain and then perform real-time sensing of neuromodulators at those neurons. For example, green SPOTIT and the red real-time dopamine sensor, RdLight1^14^ can be used together to detect real-time dopamine release in opioid-dependent neuronal circuits. Lastly, SPOTcal can label the neuronal populations relevant in opioid signaling and allow calcium imaging within those neuronal populations to further study opioid-dependent neural activity.

Overall, SPOTall and SPOTIT provide a new single-chain fluorescent integrator platform for engineering a variety of GPCR integrators. They allow mapping of GPCR agonists globally, which is important for an unbiased search of endogenous agonist release and for studying drug localization.

## Acknowledgments

Kayla Kroning is supported by F31MH12915001. Noam Gannot is supported by F31HL165733-01A1. Research is supported by R01DA05320001A1, HL156989 and AT011652.

## Author contributions

Kayla Kroning and Wenjing Wang conceived the sensor designs and experimental designs for HEK293T cell testing. Peng Li conceived the experimental designs for animal testing. Kayla Kroning performed experiments for Fig. 1, Fig. 2, Fig. 3a and 3b, Supplementary Fig. 1, and Supplementary Fig. 8. Noam Gannot determined the animal testing protocol and performed the experiments for Fig. 4 and Supplementary Fig. 5. Xingyu Li performed the experiment for Fig. 5 and Supplementary Fig. 9. Aubrey Putansu performed the experiment for Supplementary Fig. 2. Jennifer Sescil performed the experiment for Supplementary Fig. 6 and Supplementary Fig. 7. Jiaqi Shen performed the experiment for Supplementary Fig. 3. Guanwei Zhou performed the experiments for Fig. 3c and 3d and Supplementary Fig. 4. Hailey Fiel and Avery Wilson contributed to cloning of the constructs used in these experiments. All authors participated in manuscript writing and editing.

## Data Availability

All the data supporting the findings in this manuscript are supplied within the main manuscript and the supplementary information files. All the DNA constructs used in this study are available upon request to the corresponding author.

## Methods

### Plasmids, cloning, and virus preparation

Constructs were cloned in an ampicillin-resistant adeno-associated viral vector with a CAG promoter. Standard cloning procedures, such as Q5 or Taq polymerase PCR amplification, NEB restriction enzyme digest, and T4 ligation or Gibson assembly were used. Heat shock transformation with XL1-blue competent cells was used for transformation of plasmids. To prepare concentrated AAV1/2 virus, the protocol described in Shen et al. was used.

<u>Reference:</u> Shen, J. *et al.* A general method for chemogenetic control of peptide function. *Nature Methods* **20**, 112-122 (2023).

### HEK293T cell culture and transfection

Complete growth media was used for HEK293T cell culture. Complete growth media includes: 1:1 DMEM (Dublecco’s Modified Eagle medium, GIBCO): MEM (Modified Eagle medium, GIBCO), 10% FBS (Fetal Bovine Serum, Sigma), 1% (v/v) penicillin-streptomycin (Gibco).

Cells were plated at a density so that they would reach 80-90% confluence on the day of transfection. Cells were cultured at 37 °C under 5% CO_2_.

For transfection, we pre-treated 48-well plastic plates with 200 μL of 20 μg/mL human fibronectin (Milipore Sigma) for 10 minutes at 37 °C under 5% CO_2_. After 10 minutes, we removed the fibronectin and added 200 μL of 80-90% confluent HEK293T cells into each well. To prepare the transfection mixture for each well, we combined 100 ng of DNA with 1 μL of PEI MAX solution (polyethyleneimine, Polysciences) in 10 μL of DMEM and incubated at room temperature for 10 minutes. After the incubation, we added 100 μL of complete growth media to the DNA-PEI MAX mixture and mix it with the cell suspension in the well. Cells were incubated at 37 °C with 5% CO_2_ until stimulation 20-24 hours later.

### HEK293T cell fixation and immunostaining

Cells were removed and 100 μL of 4% formaldehyde was added per well. Formaldehyde was incubated for 20 minutes at room temperature and then removed. The wells were then washed twice with phosphate buffer saline (PBS). After fixation, the cells were permeabilized using 100 μL of prechilled methanol per well and incubated at -20 °C for 5 minutes. The wells were then washed twice with PBS. For immunostaining, 1:1000 chicken anti-GFP antibody (primary antibody) and 1:1000 anti-chicken-647 (secondary antibody) were diluted in PBS with 1% BSA. 100 μL of primary antibody was added to each well and then the plate was rocked at room temperature for 30 minutes. After 30 minutes, the cells were washed twice with PBS and antibody staining was repeated with the secondary antibody. After 30 minutes of secondary antibody rocking, the cells were washed twice with PBS and then fixed with 4% formaldehyde again for 20 minutes at room temperature. Adparafter 20 minutes, the cells were washed 2x with PBS and then 200 μL of pH 11 CAPS buffer was added to the cells, except for the cells for the pH titration. For experiments with red fluorescent proteins, mouse anti-FLAG and anti-mouse-647 primary and secondary antibodies, respectively, were used following the same dilutions and procedure.

### Confocal microscopy of HEK293T cells

Confocal imaging was performed on a Nikon inverted confocal microscope with 20x air objective and 60x oil immersion objective, outfitted with a Yokogawa CSU-X1 5000RPM spinning disk confocal head, and Ti2-ND-P perfect focus system 4, a compact 4-line laser source: 405 nm (100 mW) 488 nm (100 mW), 561 nm (100mW) and 640-nm (75 mW) lasers. The following combinations of laser excitation and emission filters were used for various fluorophores: EGFP/Alexa Fluor 488 (488 nm excitation; 525/36 emission), mCherry (568 nm excitation; 605/52 emission), Alexa Fluor 647 (647 nm excitation; 705/72 emission), and differential interference contrast (DIC). The acquisition time for all images was 1 second with 50% laser power intensities. ORCA-Flash 4.0 LT+sCMOS camera. 20x objective or 60x objective was used for all HEK293T cell experiments. 10x objective was used for mouse brain slice images. All images were collected using Nikon NIS-Elements hardware control and processed using NIS-Elements General Analysis 3 software.

### Analysis of HEK293T cell and animal images

For all HEK293T cell experiments, three technical replicates were performed (three wells for each condition.) 10 images were taken per well, and the sum intensity value was measured for each image. The sum intensity value was taken using the NIS-Elements General Analysis 3 software. Specifically, a threshold value was set to be above the autofluorescence of the cells, where only real cpGFP/cpRFP fluorescence signal would be measured. The threshold was generally set to be 1.5-2x higher than the autofluorescence from the plate void of cells. The same threshold was used for all conditions in a single experiment. Two values were taken from the cells that have fluorescence values within the threshold: the mean intensity value and the total object area of the field of view. The total object area is the area of cells that fit in the threshold and the mean intensity value is the mean intensity value of the cells counted in the total object area. The total object area was then multiplied by the mean intensity value to calculate the sum intensity value.

For background subtraction of HEK293T cell experiments, the mean intensity value of the plate by itself (area with no cells) was multiplied by the object area and then subtracted from the sum. For background subtraction of mouse experiments, the mean intensity value of the region of the brain away from the injection site was multiplied by the object area and then subtracted from the sum. For all HEK293T cell experiments and the M-SPOTIT2 mouse experiment, one background value for subtraction was used for each experiment. For mouse testing of B2AR-SPOTall, we performed different background subtractions depending on the background of each individual image. This was because the dimmer fluorescent signal of B2AR-SPOTall is more impacted by tissue autofluorescence.

For HEK293T cell experiments, the sum intensity values for the 10 images from each technical replicate were averaged and then each technical replicate was plotted as one point in the graph. For the mouse experiments, sum intensity values were calculated for each individual image. The mean of the sum intensity for the 5 brightest images from each mouse was calculated and plotted as one point in the graph. For all experiments, Prism GraphPad software was used for plotting, as well as performing unpaired two-sided Student’s t tests to calculate significance. All images were included in analysis, except for images that had lower than a 50% cell density in the image, were blurry, or had auto fluorescent artifacts. Images excluded from analysis are indicated in the data sheets included in the submission.

### B2AR-SPOTall initial testing and engineering

20-24 hours after transfection, HEK293T cells were stimulated with 100 μL of isoproterenol diluted in complete growth media, where there would be a 10 μM final concentration within the well. 24 hours after stimulation, the cells were fixed, immunostained, and imaged with a CAPS pH 11 buffer following the above protocols. 60x imaging was performed using glass bottom plates (Corning).

### B2AR-SPOTall agonist exposure experiment

20-24 hours after transfection, HEK293T cells were stimulated with 100 μL of isoproterenol diluted in complete growth media to a 10 μM final concentration within the well. Cells were incubated with isoproterenol for either 30 seconds, 5 minutes, 6 hours, and 24 hours. After the indicated exposure time, the cells were washed 2x with complete growth media and then incubated without agonist for 24 hours. The cells were then fixed and imaged at 20x with a CAPS pH 11 buffer following the above protocol.

### B2AR-SPOTall maturation experiment

HEK293T cells were transfected with B2AR-SPOTall DNA for all conditions at the same time. The 24-hour time point was stimulated 24 hours before imaging (24 hours after transfection) with 100 μL of isoproterenol diluted in complete growth media to a 50μM final concentration within the well. After 5 minutes of isoproterenol incubation, the cells were washed 2x with complete growth media. The 8, 6, 4, 3, 2, 1.5, 1, and 0.5-hour time points were stimulated 8, 6, 4, 3, 2, and 1.5 hours before imaging. After stimulation and washing, the cells were incubated without drug at 37 °C under 5% CO_2_. All cells were fixed and imaged at the same time to minimize protein level differences between conditions. Cells were imaged at 20x magnification with a CAPS pH 11 buffer.

### B2AR-SPOTall pH titration

24 hours after transfection with B2AR-SPOTall DNA, HEK293T cells were stimulated with 10μM isoproterenol for 5 minutes. The cells were then washed 2x with complete growth media and incubated for 24 hours in complete growth media without drug at 37 °C under 5% CO_2_. The cells were then fixed and imaged with buffers of different pH. pHs: 5.61, 6.96, 7.66, 8.06, 8.62, 9.12, 10.25, and 11.01. 100 mM Tris-HCl, 100 mM CAPS, and 1x PBS buffers were used. Cells were imaged at 20x magnification.

### B2AR-SPOTall selectivity

24 hours after transfection with B2AR-SPOTall DNA, HEK293T cells were stimulated with different agonists to give a 50 μM final concentration. The agonists were diluted in complete growth media and added for 5 minutes. After 5 minutes of agonist incubation, the wells were washed 2x with complete growth media, and the cells were incubated without drug at 37 °C under 5% CO_2_. 24 hours after stimulation, the cells were fixed and imaged at 20x magnification with a CAPS pH 11 buffer.

### CHRM2- and DRD1-SPOTall initial testing

HEK293T cells expressing the CHRM2- or DRD1-SPOTall sensors were stimulated with iperoxo and dopamine, respectively to a final concentration of 100 μM. After stimulation, the cells were incubated at 37 °C under 5% CO_2_ for 24 hours. Then, the cells were fixed, immunostained, and imaged at both 60x and 20x magnifications with a CAPS pH 11 buffer. Glass bottom plates were used.

### M- and K- red-SPOTIT initial testing and engineering

HEK293T cells expressing the M- and K-red SPOTIT sensors were stimulated with fentanyl or salvinorin A, respectively, to a final concentration of 10 μM. After stimulation, the cells were incubated at 37 °C under 5% CO_2_ for 24 hours. Then, the cells were fixed, immunostained, and imaged at both 60x and 20x magnifications with a CAPS pH 11 buffer. Glass bottom plates were used.

### B2AR-SPOTall agonist titrations

HEK293T cells expressing B2AR-SPOTall were stimulated with varying concentrations of the following drugs: epinephrine, isoproterenol, arformoterol, indacaterol, levalbuterol, and butoxamine. Serial dilutions were performed, where the drug was diluted in complete growth media. Drug was added to the well for 5 minutes and then the wells were washed 2x with complete growth media. After stimulation, the cells were incubated at 37 °C under 5% CO_2_ for 24 hours without drug. Then, the cells were fixed and imaged at 20x magnification with a CAPS pH 11 buffer.

### Calcium-Gated SPOTIT

HEK293T cells expressing SPOTcal were stimulated with fentanyl diluted in complete growth media to a final concentration of 10 μM. After stimulation, the cells were incubated at 37 °C under 5% CO_2_ for 18 hours. For calcium and ionomycin stimulation, 100 μL of ionomycin and CaCl_2_ diluted in pre-warmed complete growth media were added to the wells to a final concentration of 2 μM and 5 mM, respectively. Images were taken every 1 second for 2 minutes immediately before and after calcium stimulation. 20x magnification imaging was used.

### Gai binding site mutations testing

#### cAMP Assay

100 ng of SPOTIT DNA and 50ng of GloSensor DNA (Promega) was used for transfection. 20 hours after transfection, the complete growth media in the well was replaced with 100 µL of 2 mM D-luciferin potassium salt (Gold Bio, #LUCK) in complete growth medium (with 50 mM HEPES). Luminescence values were measured using a plate reader (BioTek, CYTATION 5). Luminescence values were measured for 1 hour, after which cells were treated with 1 µL of 100 µM forskolin (Sigma-Aldrich, #F6886) in complete growth media. Luminescence values were measured for 30 min, and then the opioid condition cells were treated with 1 µL of 1 mM fentanyl (for mu-opioid receptor constructs) or 1 µL of 1 mM Sal A (for kappa-opioid receptor constructs) in complete growth media. Luminescence values were measured for 30 min.

#### Confocal Microscopy

24 hours after transfection of the SPOTIT sensors, HEK293T cells were stimulated with 100 µL of salvinorin A (for kappa-opioid receptor constructs) or fentanyl (for mu-opioid receptor constructs) diluted in complete growth media to a final concentration of 10 µM. After stimulation, the cells were incubated at 37 °C for 20 hours and then fixed and imaged at 20x with a CAPS pH 11 buffer.

### Mouse Lines

All procedures were carried out in accordance with the animal care standards in National Institutes of Health (NIH) guidelines and approved by the University Committee on Use and Care of Animals at the University of Michigan. Animals were maintained on a 12 hours light/dark cycle with ad libitum access to food and water. All mice strains utilized were C57BL/6J genetic background.

### M-SPOTIT2 mouse testing

*Stereotactic injection.* Adult mice were anesthetized with isoflurane (4 - 5% for induction, 1.5% for maintenance) for stereotactic injection. Animals received 5 mg/kg of preemptive analgesic carprofen prior to placement in the stereotactic frame (David Kopf Instruments) and body temperature was maintained at 35 – 37 °C using a feedback-controlled heating pad (Physitemp, TCAT-2LV). The following coordinates were used for injection: PreBotC, ± 1.3 mm from the midline; - 6.7 mm posterior to the bregma, −6.1 mm ventral from bregma. 500nL of the AAV1/2-CAG-M-SPOTIT2 virus was injected into the PreBotC at a rate of 50nL/min.

#### Opioid administration and histology

6 days after injection of the viral vectors into the mouse brain, 100mg/kg of morphine or saline was administered through intraperitoneal injection. Twenty-four hours after drug or saline administration, the mice were deeply anesthetized and transcardially perfused with 10 mL PBS, followed by 10 mL 4% paraformaldehyde (PFA). The brain tissue was removed and post-fixed overnight at 4 °C in 4 % PFA and then cryopreserved in 30% sucrose for 48 hours. The processed tissue was then embedded into optimum cutting temperature compound (OCT) and sectioned into serial slices through the relevant region at 40 microns with a Leica cryostat CM3050 S cryostat.

For immunostaining, sections were washed in 0.1% PBS Tween 20 (PBS with 0.1% Tween 20) and permeabilized with 0.3% PBS Triton X-100 for 30 min, followed by 2% bovine serum albumin (BSA) blocking solution at room temperature for 1 hour. Sections were then incubated with primary antibody (Chk pAb to GFP; Abcam13970; 1:500 dilution; Lot: GR3361051-18) in BSA overnight at 4 °C. After washing in 0.1% PBS Tween 20 three times (5 minutes each time), sections were incubated with secondary antibody (Alexa Fluor 647 goat anti-chicken IgG; Life technologies A21449; 1:500 dilution; Lot: 1599393) for 2 hrs. After washing in 0.1% PBS Tween 20 three times (5 min each time), sections were then stained with 4’,6 -diamidino-2-phenylindole, dihydrochloride (DAPI; Invitrogen; D1306) diluted 1:10000 in PBS. After washing in 0.1% PBS Tween 20 three times (5 minutes each time), sections were imaged on a confocal microscope with pH 11 buffer.

### B2AR-SPOTall mouse testing

#### Stereotactic injection of AAV into the mouse brain

The procedure of stereotactic injection was described in previous work. Adult mice were anesthetized with isoflurane (5% for induction, 1.5% for maintenance), administered 5 mg kg^−1^ of carprofen, prior to being placed in a stereotactic apparatus. The mice’s shaved heads were disinfected using one application of betadine and three applications of alcohol pad. The body temperature of the was monitored and maintained at 35 °C. A total of 400 nl of concentrated AAV1/2-CAG-B2AR-SPOTall were injected into the lateral hypothalamic area (0.95 mm lateral to midline, −1.40 mm posterior, and −5.25 mm ventral to bregma) at a rate of 50nl min^−1^ for 8 minutes. After injection, the pipette was left in the brain for 10 minutes to allow for pressure to equalize.

#### Isoproterenol, epinephrine administration and histology

After the injection of the viral vectors into mouse brain, a local injection of 1 μl 10 mM Isoproterenol, epinephrine, or saline (Hospira, #00409-4888-10) control was administered into LHA via stereotactic injection seven days later. Twenty-four hours after local injection of drugs or saline, the animals were euthanized and perfused with PBS and 4% paraformaldehyde (PFA, Electron Microscopy Science, #15713). The brain tissues were collected and post-fixed overnight in 4% PFA then cryoprotected in 30% sucrose for 48 hours at 4 °C. The fixed tissue was then embedded in optimum cutting temperature compound, sectioned at 40 µm, and mounted onto glass slides. To prepare the tissue sections for immunostaining, tissue sections were first incubated in 0.3% Triton (Sigma, #T8787) in PBS for 30 minutes, followed by blocked in 2% BSA (Sigma, #A9647) in 0.1% Tween-20 in PBS for 1 hour. Next, the sections were incubated with chicken anti-GFP antibody (1:500, abcam, #ab13970) in 2% BSA overnight at 4 °C. After three washes with 0.1% Tween-20 for 5 minutes each, the sections were incubated with donkey anti-chicken Alexa Fluor 647 antibody (1:500, Invitrogen, #A78952) for 2 hours at room temperature. The tissue sections were then rinsed in 0.1% PBS Tween-20 and stained with DAPI (1:10,000, Invitrogen, #D1306) for 10 minutes at room temperature, and fixed with pH 11 PBS for 20 minutes. Finally, Confocal images were captured using a Nikon A1 Confocal microscope with pH 11 buffer.

### Respiratory Recording

Morphine was prepared in sterile saline at a concentration at 10 mg/mL and delivered at a dose of 100 mg/kg. To test the respiratory depression of morphine, individual mice were placed in a 450 mL whole-body plethysmography (WBP) chamber at room temperature (22°C). Mice were allowed to acclimate to the chamber for 30 minutes before the respiratory parameters were recorded by Emka IOX2 software (EMKA Technologies, Paris, France). Animals then underwent a 15 minutes baseline recording before a 200 uL IP injection of sterile saline. Following the injection, mice underwent an additional 30 minutes recording and then returned to their home cage for 20 minutes. Mice were then returned to the WBP for a 15-minutes baseline recording before an IP injection of morphine. Following the injection, the mice were recorded for an additional 30 minutes.

