## Supplementary Information for "Single-chain fluorescent integrators for mapping G-protein-coupled receptor agonists"

**This PDF file includes:**

Figures S1 to S9
Legends for Movies S1 to S4

**Other supporting materials for this manuscript include the following:**

Movies S1 to S4

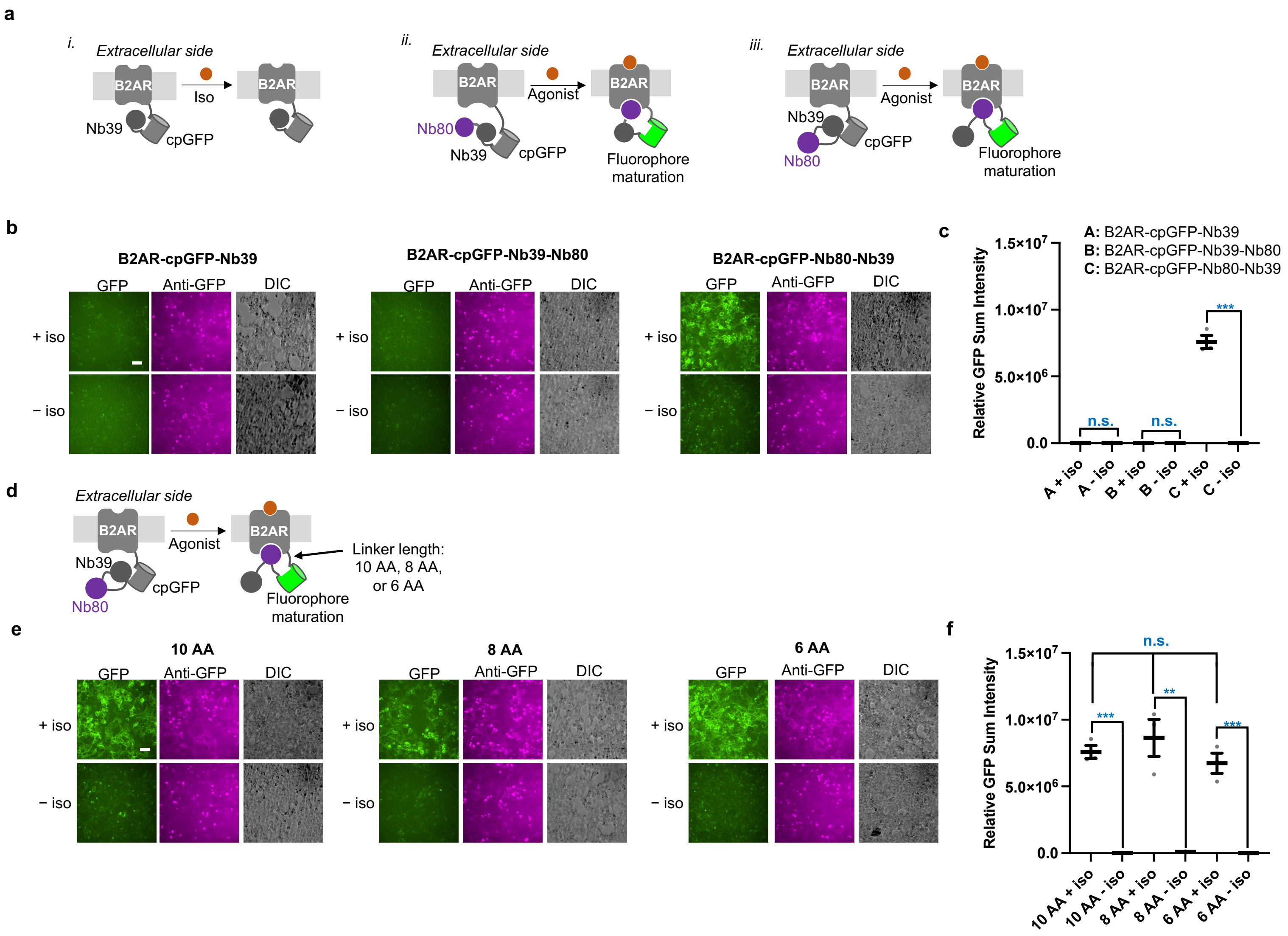


**Fig. S1. Engineering of B2AR-SPOTall**. **a**, Schematic of the different B2AR-SPOTall geometries we tested. **b**, HEK293T cell testing of the different geometries shown in **a**. HEK293T cells expressing the B2AR-SPOTall sensors were stimulated with 10 μM isoproterenol (iso). **c**, Quantification of **b**. A: n.s. p= 0.6278, B: n.s. p= 0.7513, C: ***p= 0.0001. *n*= 3. **d**, Schematic indicating what linker region we truncated for the B2AR-SPOTall sensor. **e**, HEK293T cells expressing the B2AR-SPOTall sensors were stimulated with 10 μM isoproterenol. **f**, Quantification of **e**. 10AA: ***p= 0.001, 8 AA: **p= 0.0036, 6 AA: ***p= 0.0009, comparing + iso conditions for 10 AA, 8 AA, and 6 AA: n.s. p= 0.5101, 0.2953, 0.4046. *n*= 3. For **b** and **e**, cells were fixed, immunostained, and imaged at pH 11 24 hours post stimulation. GFP, cpGFP fluorescence. Anti-GFP, protein expression level. DIC, differential interference contrast. Scale bar, 20 μm. For **c** and **f**, Error bars are the SEM. The thick horizontal bar is the mean value of three technical replicates. The number above the dots is S/N and the stars indicate significance compared to the "-iso" conditions. Significance was calculated using an unpaired, two-tailed Student's *t*-test. A biological replicate was performed for all experiments.

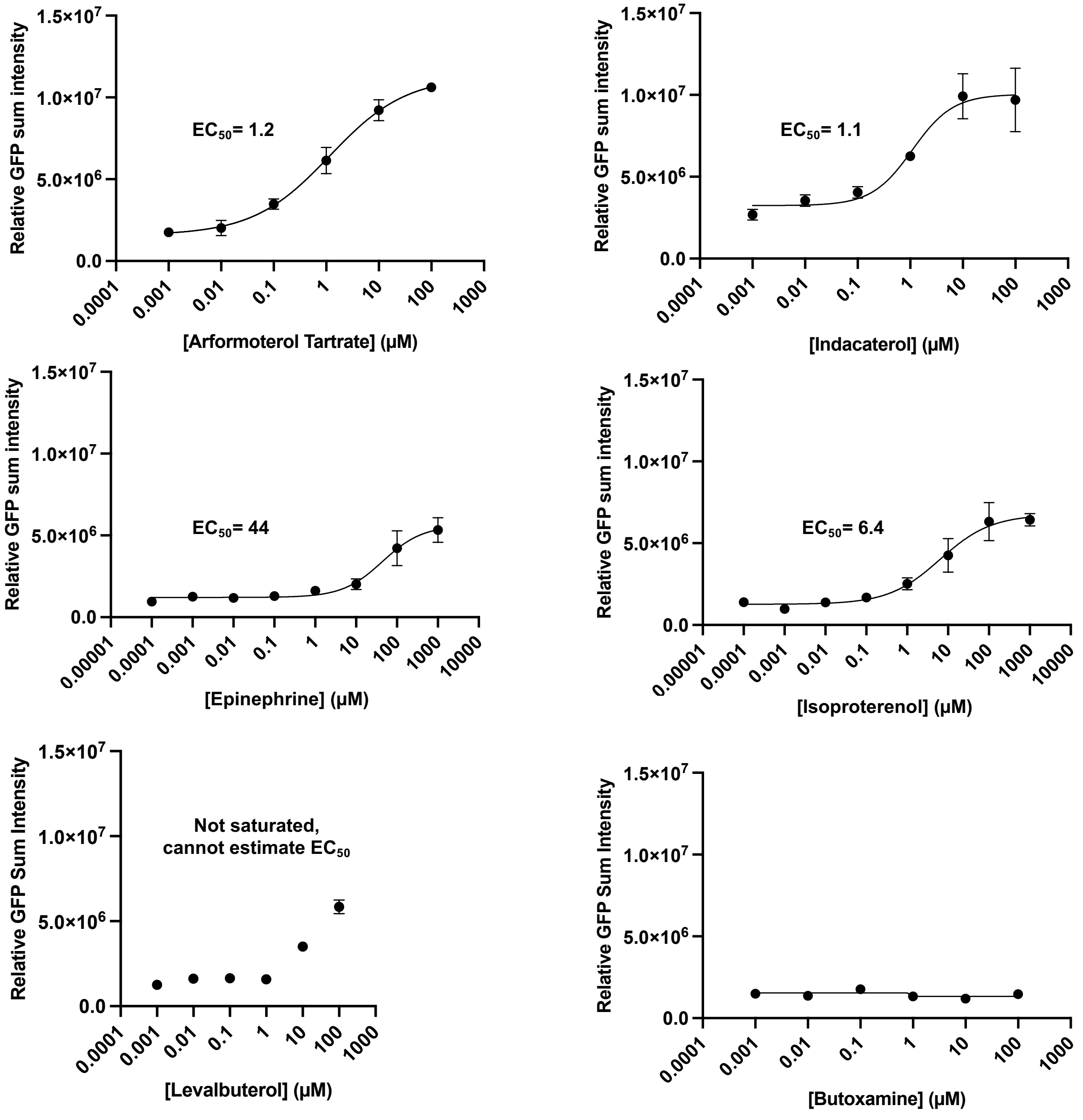


**Fig. S2. Dose response curves of the B2AR-SPOTall against a variety of ligands.** HEK293T cells expressing the B2AR-SPOTall sensor were stimulated with differing concentrations of the indicated ligand. 24 hours post stimulation, the cells were fixed and imaged at pH 11. The mean of each concentration's fluorescence response is indicated by the black dot. EC50 values are in μM. Error bars are SEM. *n*= 3.

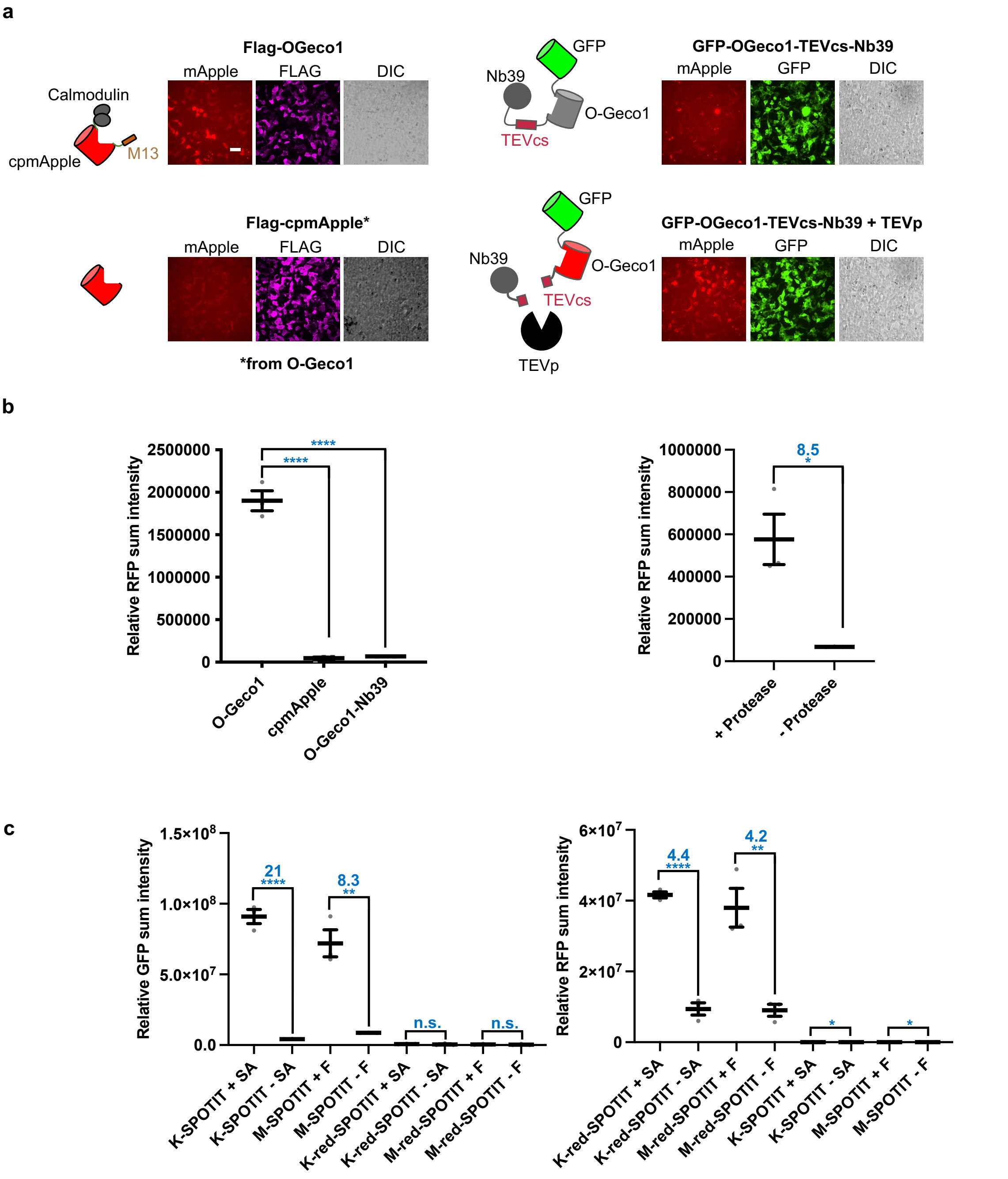


**Fig. S3. Engineering of red-SPOTIT and red-SPOTIT statistics. a**, Schematics and HEK293T cell images of red-SPOTIT engineering. HEK293T cells were fixed, immunostained, and imaged 24 hours after transfection at pH 11. mApple, cpmApple fluorescence. FLAG, protein expression levels. GFP, protein expression levels. DIC, differential interference contrast. Scale bar, 20 μm. **b**, Quantification of **a**. For the plot, error bars are the SEM. The thick horizontal bar is the mean value of three technical replicates. The number above the dots is S/N and the stars indicate significance calculated using an unpaired, two-tailed Student's *t*-test. O-Geco1-Nb39: ****p< 0.0001, cpmApple: ****p< 0.0001. Protease: *p= 0.0129. *n*= 3. **c**, Plot of red-SPOTIT drug stimulation. SPOTIT expressing cells were stimulated with 10 μM of fentanyl (F) or salvinorin A

(SA) for MOR or KOR, respectively. 24 hours post opioid stimulation, cells were fixed, immunostained, and imaged at pH 11. Error bars are the SEM. The thick horizontal bar is the mean value of three technical replicates. The number above the dots is S/N and the stars indicate significance from the “- SA” or “-F” conditions. Significance was calculated using an unpaired, two-tailed Student’s *t*-test. all p****value < 0.0001. M-SPOTIT GFP channel: **p= 0.0027. K-red SPOTIT GFP channel: n.s. p= 0.4706, M-red-SPOTIT: n.s. p= 0.4147, M-red-SPOTIT red channel: **p= 0.0072, K-SPOTIT red channel: *p= 0.0331, M-SPOTIT red channel: *p= 0.0383. *n*= 3. A biological replicate has been performed for experiments b and c that yielded similar results.

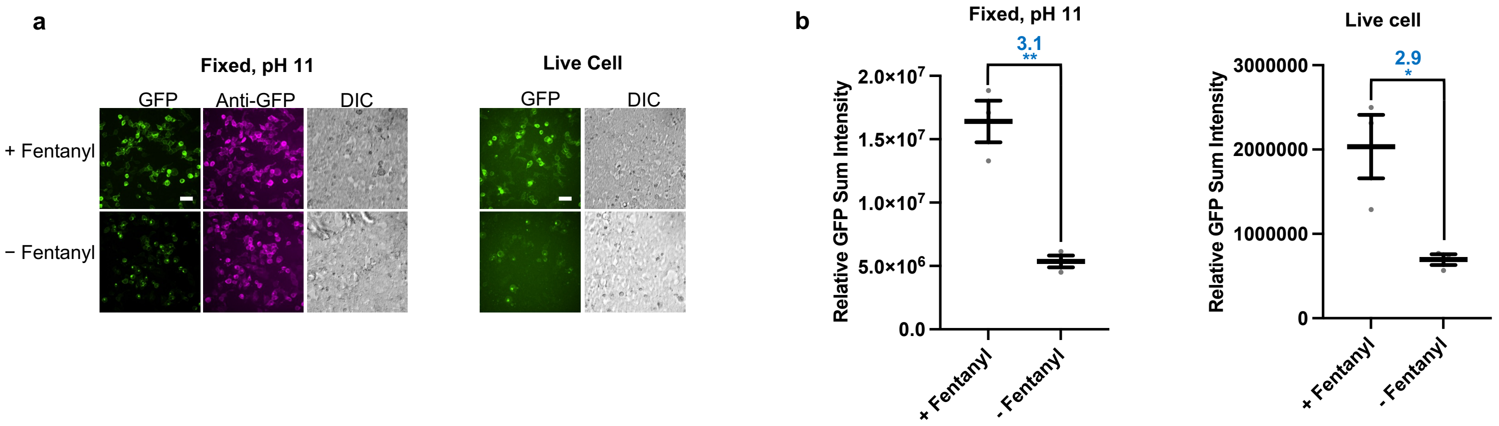


**Fig. S4. SPOTcal opioid dependence testing. a,** Opioid dependence of the opioid-activated calcium sensor in fixed cells and live cells. HEK293T cells expressing the sensor were stimulated with 10 μM fentanyl. 24 hours post stimulation, cells were fixed, immunostained, and imaged at pH 11 or were imaged live. GFP, cpGFP fluorescence. Anti-GFP, protein expression level. DIC, differential interference contrast. Scale bar, 20 μm. **b**, Quantification of **a**. For the plot, error bars are the SEM. The thick horizontal bar is the mean value of three technical replicates. The number above the dots is S/N and the stars indicate significance calculated using an unpaired, two-tailed Student's *t*-test. Live cell: *p= 0.0247, Fixed: **p= 0.0029.

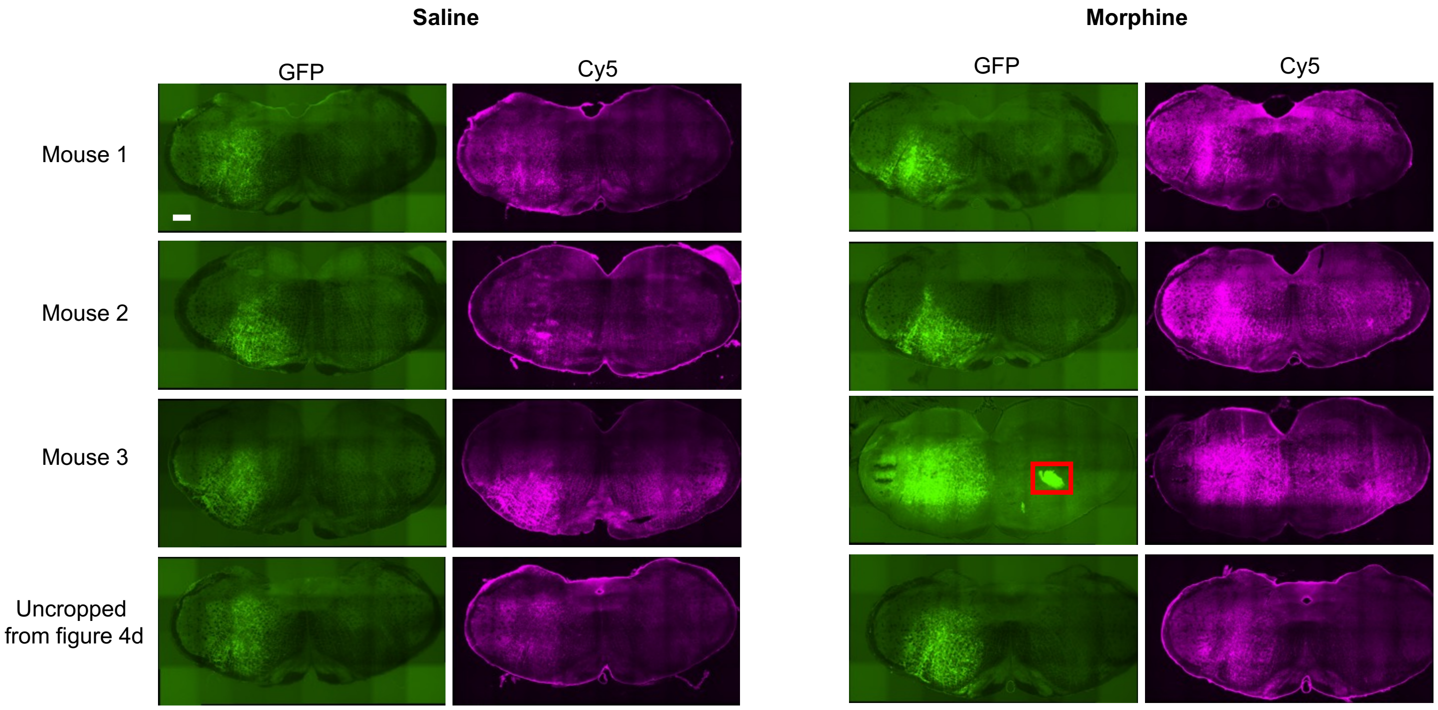


**Fig. S5. Representative images for the animal experiment shown in Fig. 4.** GFP, cpGFP fluorescence. Anti-GFP, protein levels. Red box: artifact removed for analysis. Scale bar, 300 μm.

**a**

MDSSAAPTNASNCTDALAYSSCSPAPSPGSWVNLSHLDGNLSDPCGPNRTDLGGRDSLCPPTGSPSMITAITIMALYSIVCVVGLFGNFLVMYVIVRYTKMKTATNIYIFNLALADALATST
LPFQSVNYLMGTWPFGTILCKIVISIDYYNMFTSIFTLCTMSVDRYIA**V**CH**PV**KAL**D**FRTPRNAKIINVCNWILSSAIGLPVMFMATTKYRQGSIDCTLTFSHPTWYWENLLKICVFIFAFIMPVL
IITVCYGL**M**ILRLKSVRMLSGSKE**K**DRNLR**R**ITRMVLVVVAVFIVCWTPIHIYVIIKALVTIPETTFQTVSWHFCIALGYTNSCLNPVLYAFLDENFKRCFREFCIPTSSNIEQQNSTRIRQNTRD
HPSTANTVDRTNHQLENLEAETAPLP

**b**

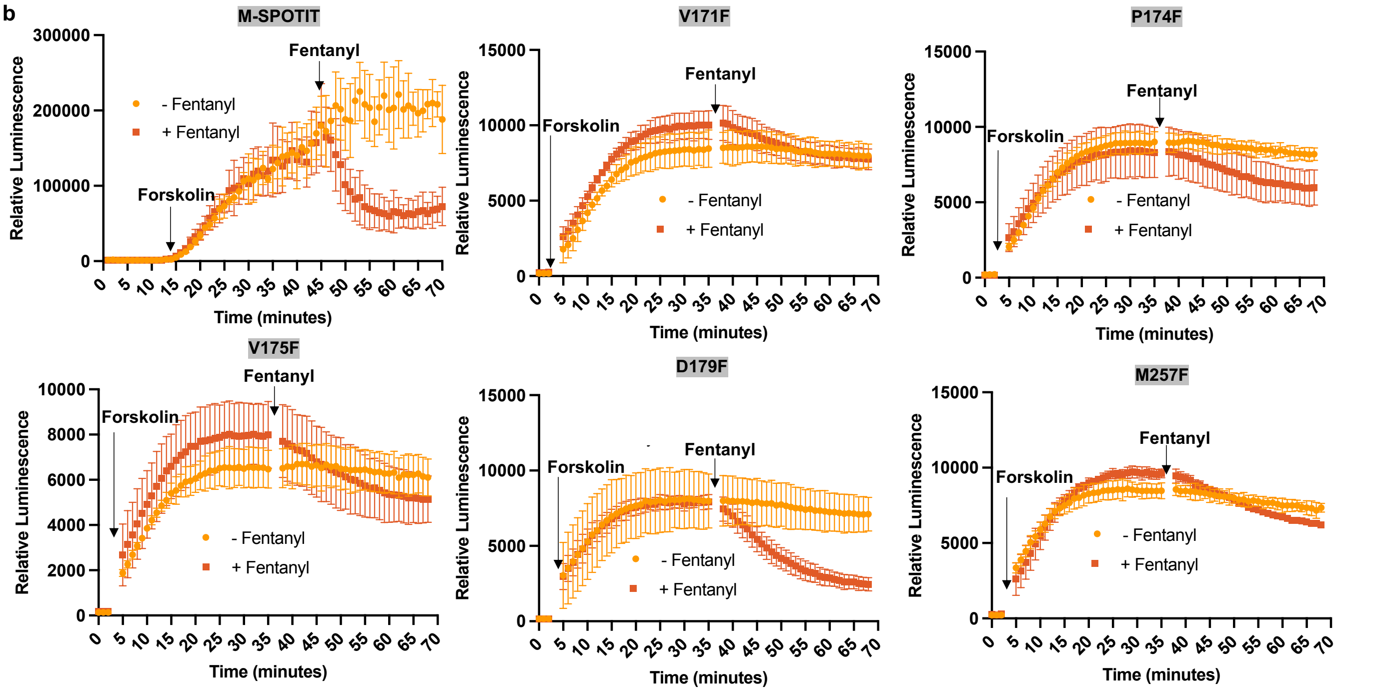


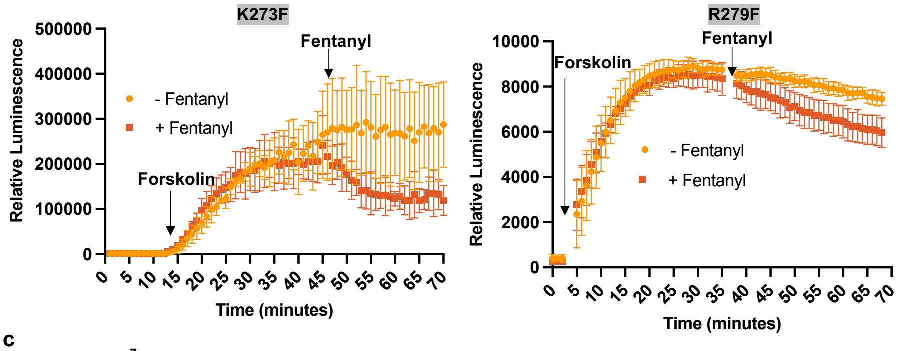


**c**

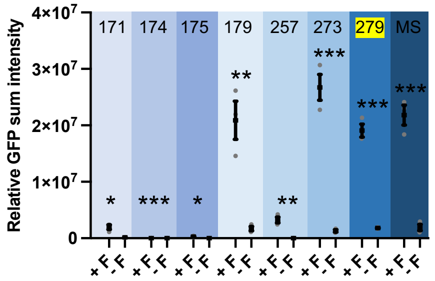


**d**

| Construct | -Fent/+Fent (Lum at 68min) | S/N (Fluorescence) |
|---|---|---|
| M-SPOTIT | 2.61 | 12 |
| V171F | 1.03 | 12 |
| P174F | 1.37 | 8400/0 |
| V175F | 1.19 | 82 |
| D179F | 2.88 | 12 |
| M257F | 1.18 | 220 |
| K273F | 2.27 | 21 |
| R279F | 1.25 | 11 |

**Fig. S6. Mutating the G-protein binding sites of M-SPOTIT. a,** MOR amino acid sequence. We mutated the residues in red to phenylalanine to decrease G-protein signaling. **b**, GloSensor assay of the different M-SPOTIT mutants to measure G-protein binding. HEK293T cells expressing the M-SPOTIT mutants and the GloSensor were stimulated with forskolin to increase cAMP levels, then stimulated with fentanyl. A decrease in luminescence indicates decreased cAMP. *n= 3.* **c**, Plot of the opioid-induced fluorescence change of the M-SPOTIT mutants. Error bars are the SEM. The thick horizontal bar is the mean value of three technical replicates. The stars indicate significance compared to the "-F" conditions. Significance was calculated using an unpaired, two-tailed Student's *t*-test. F, fentanyl. 171: *p= 0.0132, 174: ****p<0.0001, 175: *p= 0.0235, 179: **p= 0.0049, 257: **p= 0.0033, 273: ***p= 0.0004, 279: ***p= 0.0001, MS: ***p= 0.0004. *n*= 3. **d**, Summary chart of experiment. The chart includes the difference in the + Fentanyl and - Fentanyl conditions 68 minutes into measuring luminescence signal. Values closer to 1 indicate less difference. The chart also includes the fluorescence S/N of the mutants from 10 μM fentanyl stimulation. Higher fluorescence signal comparable to the original MS indicates Nb39 can still bind to the MOR.

**a**

MDSPIQIFRGEPGPTCAPSACLPPNSSAWFPGWAEPDSNGSAGSEDAQLEPAHISPAIPVIITAVYSVVFVVGLVGNSLVMFVIIRYTKMKTATNIYIFNLALADALVTTTMPFQ
STVYLMNSWPFGDVLCKIVISIDYYNMFTSIFTLTMMSVDRYIAVCHPVKALDFRTPLKAKIINICIWLLSSSVGISAIVLGGTKVREDVDVIECSLQFPDDDYSWWDLFMKICVFI
FAFVIPVLIIIVCYTLMILRLKSVRLLSGSREKDRNLR**R**ITRLVLVVVAVFVVCWTPIHIFILVEALGSTSHSTAALSSYYFCIALGYTNSSLNPILYAFLDENFKRCFRDFCFPLKMR
MERQSTSRVRNTVQDPAYLRDIDGMNKPVQ

**b**

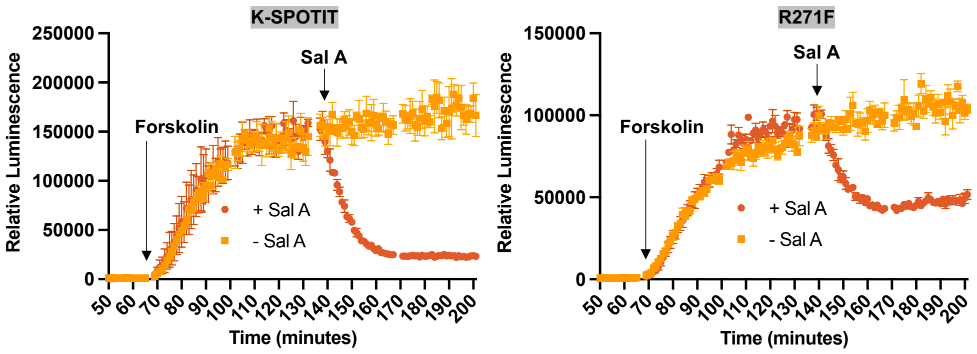


**d**

| Construct | -SA/+SA (Lum at 68min) | S/N (Fluorescence) |
|---|---|---|
| K-SPOTIT | 7.19 | 34 |
| R271F | 2.01 | 330 |

**c**

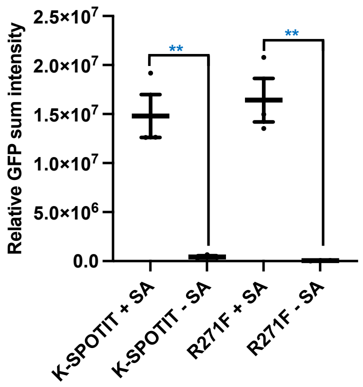


**Fig. S7. Mutating the G-protein binding sites of K-SPOTIT. a,** KOR amino acid sequence. We mutated the residue in red to phenylalanine to decrease G-protein signaling. **b**, GloSensor assay of the K-SPOTIT mutant to measure G-protein signaling. HEK293T cells expressing the K-SPOTIT mutant and the GloSensor were stimulated with forskolin to increase cAMP levels, then stimulated with salvinorin A. A decrease in luminescence indicates decreased cAMP. *n= 3.* **c**, Plot of the opioid-induced fluorescence change of the K-SPOTIT mutant. For the plot, error bars are the SEM. The thick horizontal bar is the mean value of three technical replicates. Significance was calculated using an unpaired, two-tailed Student's *t*-test. SA, salvinorin A. **p values: 0.0018 and 0.0028. *n*= 3. **d**, Summary chart of experiment. The chart includes the difference in the + SA and - SA conditions 68 minutes into measuring luminescence signal. Values closer to 1 indicate less difference. The chart also includes the fluorescence S/N of the mutants from 10 μM salvinorin A stimulation. High fluorescence signal comparable to the original K-SPOTIT indicates Nb39 can still bind to the KOR.

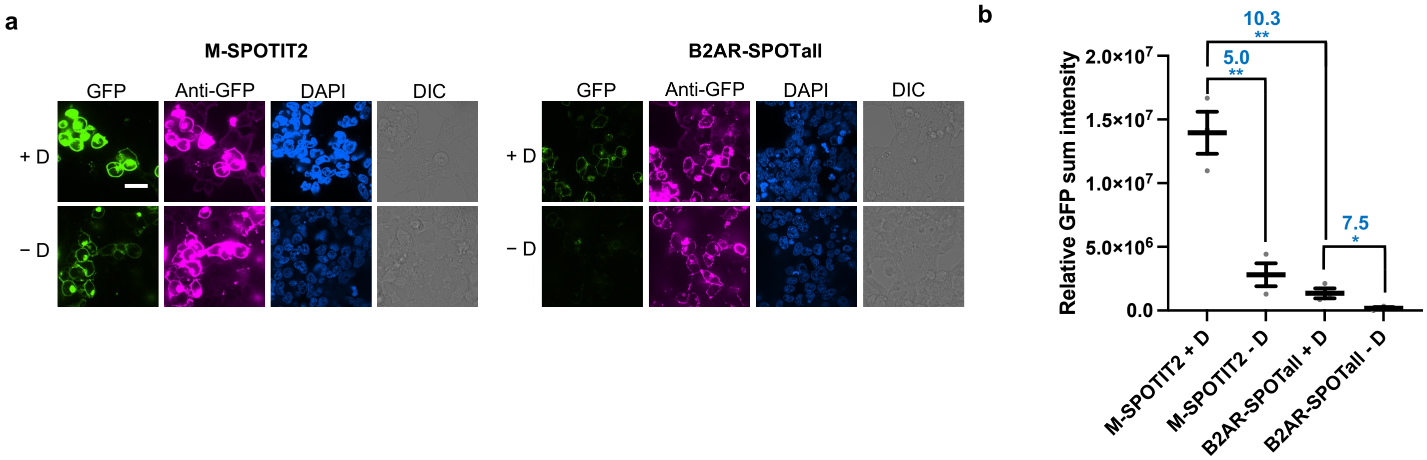


**Fig. S8. Comparing the brightness of agonist-stimulated M-SPOTIT2 and B2AR-SPOTall. a,** HEK293T cells expressing the B2AR-SPOTall or M-SPOTIT2 sensors were stimulated with 10 μM of isoproterenol or fentanyl, respectively. 24 hours post stimulation, the cells were fixed and imaged at pH 11. GFP, cpGFP fluorescence. Anti-GFP, protein expression. DAPI, nuclear staining. DIC, differential interference contrast. Scale bar, 20 μm. **b**, Quantification of experiment described in **a**. The thick horizontal bar is the mean value of three technical replicates. Error bars are SEM. The number above the dots is S/N and the stars indicate significance. Significance was calculated using an unpaired, two-tailed Student's *t*-test. **p= 0.0041 and 0.0018, *p= 0.0444. *n*= 3. A biological replicate was performed for this experiment that yielded similar results. D, drug.

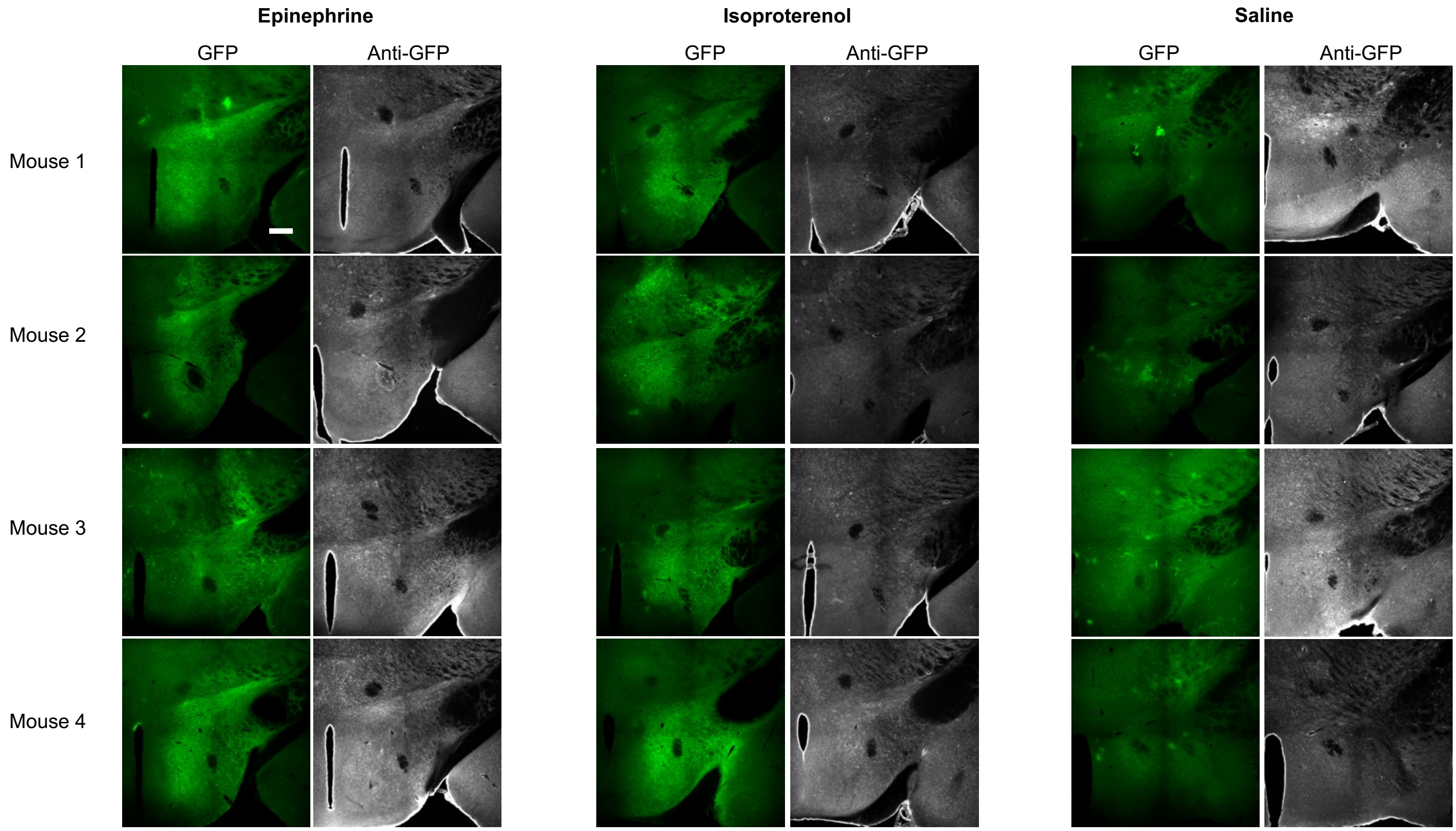


**Fig. S9. Representative images for the animal experiment shown in Fig. 5.** GFP, cpGFP fluorescence. Anti-GFP, protein levels. Scale bar, 300 µm.

**Movie S1 (separate file). SPOTcal expressing HEK293T cells stimulated with calcium and fentanyl.** HEK293T cells expressing SPOTcal were stimulated with 10 μM fentanyl. 24 hours after fentanyl stimulation, cells were stimulated with 2 μM ionomycin and 5 mM $CaCl_2$. Cells were imaged in real-time immediately before and after ionomycin and calcium stimulation.

**Movie S2 (separate file). SPOTcal expressing HEK293T cells stimulated with calcium.** **H**EK293T cells expressing SPOTcal were stimulated with a media blank. 24 hours after stimulation with the blank, cells were stimulated with 2 μM ionomycin and 5 mM $CaCl_2$. Cells were imaged in real-time immediately before and after ionomycin and calcium stimulation.

**Movie S3 (separate file). SPOTcal expressing HEK293T cells stimulated with fentanyl.** HEK293T cells expressing SPOTcal were stimulated with 10 μM fentanyl. 24 hours after fentanyl stimulation, cells were stimulated with a media blank. Cells were imaged in real-time immediately before and after the addition of the media blank.

**Movie S4 (separate file). SPOTcal expressing HEK293T cells not stimulated.** HEK293T cells expressing SPOTcal without prior stimulation with fentanyl were imaged in real-time immediately before and after the addition of a media blank.
